# Small lakes in big landscape: External drivers of littoral ecosystem in high elevation lakes

**DOI:** 10.1101/034553

**Authors:** Dragos G. Zaharescu, Carmen I. Burghelea, Peter S. Hooda, Richard N. Lester, Antonio Palanca-Soler

**Affiliations:** Biosphere 2, University of Arizona, Tucson, Arizona, USA; School of Built and Natural Environments, Kingston University London, UK; Faculty of Biological Sciences, University of Vigo, Vigo, Spain; Formerly at Birmingham University Botanic Gardens, Birmingham, U.K. Passed away in April 2006 in Birmingham, U.K.

**Keywords:** Altitude lakes, Littoral zone, Benthic invertebrates, Scale dependency, Catchment heterogeneity, Riparian vegetation, Predation, Environmental change

## Abstract

Graphical abstract:

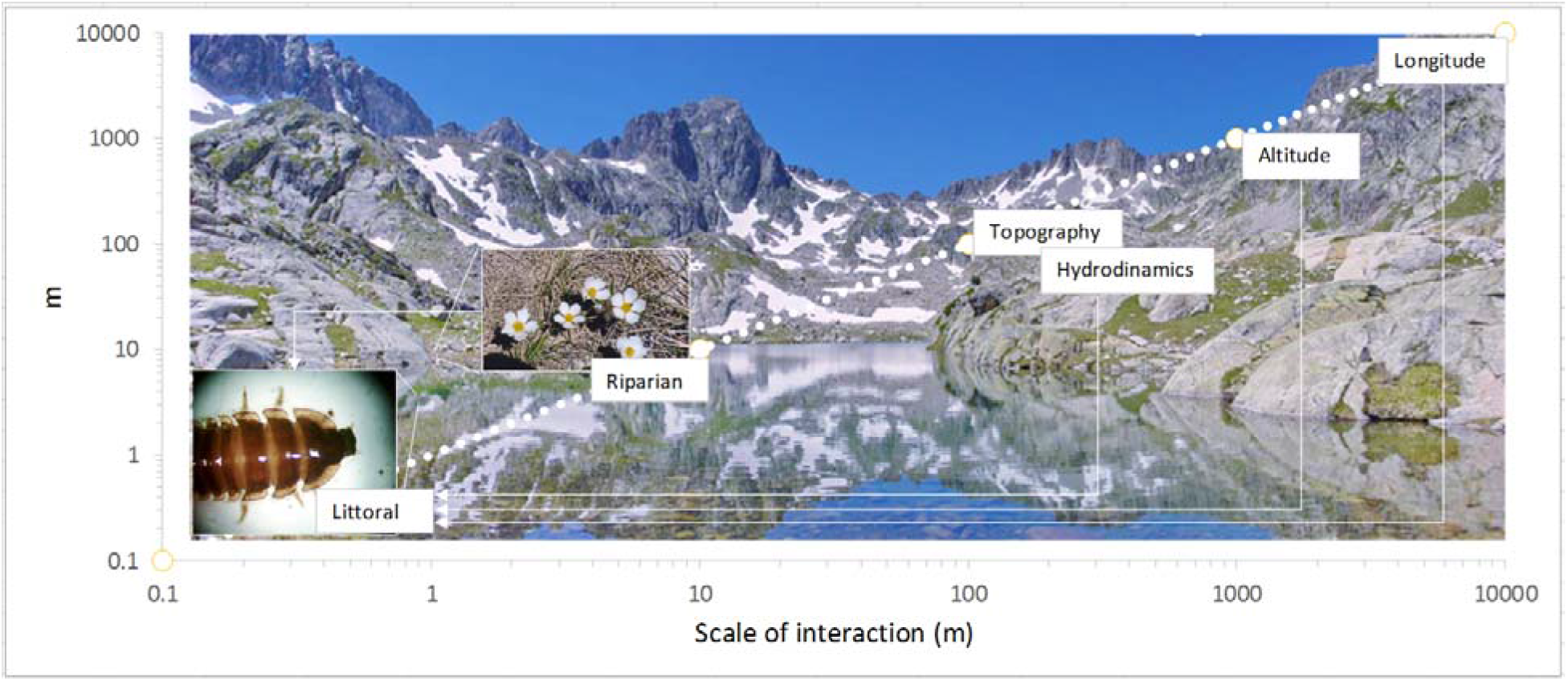

In low nutrient alpine lakes, the littoral zone is the most productive part of the ecosystem, and it is a biodiversity hotspot. It is not entirely clear how the scale and physical heterogeneity of surrounding catchment, its ecological composition, and larger landscape gradients work together to sustain littoral communities.

A total of 114 alpine lakes in the central Pyrenees were surveyed to evaluate the functional connectivity between catchment physical and ecological elements and littoral zoobenthos, and ascertain their effect on community formation. At each lake, the zoobenthic composition was assessed together with geolocation (altitude, latitude and longitude), catchment hydrodynamics, geomorphology, topography, riparian vegetation composition, the presence of trout and frogs, water pH and conductivity.

Uni- and multidimensional fuzzy set ordination models integrating benthic biota and environmental variables revealed that at geographical scale longitude surpassed altitude in its effect on littoral ecosystem, reflecting a sharp transition between Atlantic and Mediterranean bioregions. Topography (through its control of catchment type, summer snow coverage, and connectivity with other lakes) was the largest catchment-scale driver, followed by hydrodynamics (waterbody size, type and inflow/outflow volumes). Locally, riparian plant composition significantly related to littoral community structure, richness and morphotype diversity. These variables, directly and indirectly create habitats for aquatic and terrestrial stages of invertebrates, and control nutrient and water cycles. Three ecologically diverse associations characterised distinct lake sets. Vertebrate predation, water conductivity and pH (broad measures of total dissolved ions/nutrients and their bioavailability) had no major influence on littoral taxa.

The work provides exhaustive information from relatively pristine sites, which unveil a strong connection between littoral ecosystem and catchment heterogeneity at scales beyond the local environment. This underpins their role as sensors of local and large-scale environmental changes, and can be used to evaluate further impacts.

## 1 Introduction

Integrative efforts linking landscape-scale biogeochemical, hydrological and ecological processes have been intensified in the last decade, and true whole-catchment perspectives are starting to crystalize (Richter and Billings 2015). High altitude catchments are of increased relevance, partly because they are younger than the average landscape, and they are major drivers of hydrological and biogeochemical cycles affecting the wider biosphere. Their high topography, remoteness and climate allow for the formation of waterbodies of unmatched water quality, which are ecological, biogeochemical and aesthetic hotspots.

Only across Europe, there are over 50,000 remote mountain lakes (Kernan et al., 2009), of which the Pyrenees, a relatively low-density lacustric region, accounts for an estimated 4000 (Castillo-Jurado, 1992). The littoral and riparian zones of these lakes are critical mediators between sediment and nutrient fluxes from the surrounding terrestrial area and lake internal processes. Littoral surfaces also experience cross-ecosystem water and nutrient exchanges (both, autochthonous and allochthonous) with riparian zones, and provide habitat and resources for both aquatic and emerging stages of many aquatic taxa, such as most benthic insects (Gregory et al., 1991; Jonsson and Wardle; 2009; Kopacek et al., 2000). The Pyrenees are estimated to sum >797 km of littoral zone in lakes above 1000m, which are of at least 0.5ha (Castillo-Jurado, 1992), meaning that littoral processes represent a great portion of the nutrient fluxes in the catchment.

The topography, the hydrology, the bedrock geology and the climate control the intensity of bedrock weathering and nutrient transport into high altitude lakes; this influences water and sediment chemistry, and ultimately their ecosystems (Vollenweider, 1968). Even though the littoral zone is just a fraction of total lake area, it harbours the vast majority of species in a lake, and the littoral nutrient productivity is vital for aquatic food webs, contributing substantially to the whole lake ecosystem energy budget (Vander-Zanden et al., 2006, Vadeboncoeur et al., 2011).

The challenges from inhabiting shallow lake areas at high elevation range from high solar radiation and water level fluctuations, to low food availability, a short growing season, irregular freezing periods and strong seasonal temperature variation (Bretschko, 1995). Most of aquatic invertebrates are at their distributional boundaries, and they are highly sensitive to environmental changes (Bandyopadhyay et al., 1997). For example, winter mortality is a major factor regulating alpine lake macroinvertebrate populations (Oswood et al., 1991). Food availability and duration of ice/snow cover during winter are other factors affecting littoral macroinvertebrate communities (Bretschko, 1995), as there are also nitrate and ammonia levels, fish presence, lake morphology (Kernan et al., 2009) and type of shore coverage (Füreder et al., 2006).

High topography and low available nutrients generally support simple littoral ecosystems, characterized by a limited number of species and trophic levels, which are highly adapted to the local environment. Research has shown that in mountain lakes, variability in external condition can affect littoral macroinvertebrate abundances, through relative control on the proximal environment (Kernan et al., 2009). Moreover, geographical location can have a greater influence on macroinvertebrate communities than local environment (Kernan et al., 2009). It is expected that these topographical and climate restrictions introduce strong biogeographical variability, and segregation of littoral macroinvertebrates into distinct communities. Climate/environmental changes would further disrupt this natural heterogeneity, through mechanisms that alter the temperature, water and nutrient fluxes, significantly changing lake ecosystem balances. For example alpine stream benthic invertebrate communities can be particularly sensitive to climate change-driven glacier retreat (Khamis et al., 2014).

Despite a great ecological and geochemical importance of the alpine lakes littoral zone, the scale and complexity of its connectivity to surrounding landscape remains an open question. To better anticipate its response to environmental changes it is, therefore, imperative to integrate the littoral surfaces into the mechanistic understanding of how physical and ecological heterogeneity of the catchment and littoral ecosystem interact across scales before major alterations occur. This study attempts to evaluate the magnitude of influence catchment attributes have on macrozoobenthos community composition at scales from a lake to large geographical gradients. And, to assess how these interactions determine the formation of littoral associations, which can potentially serve as sensors of environmental change. We hypothesize that while local littoral environment directly sustains the macroinvertebrate community, its composition is sensitive to landscape processes at scales beyond that of the lake, through mechanisms that can affect both, aquatic and terrestrial phases of its taxa. The study area has the advantages of being at the confluence of four major biogeographical regions: Atlantic, Continental, Mediterranean, and Alpine, which should facilitate capturing the large-scale heterogeneity in a relatively narrow region.

## 2 Methodology

### 2.1 The lakes under study

A total of 114 lakes and ponds were surveyed in July 2001 in the axial Pyrenees, between degrees: 42°51'34.76" – 42°43'8.19"N and 0°29'44.39"W – 0° 8'40.29"E (Fig. 1, Supplementary List 1). Their selection was largely dictated by their accessibility, and comprised a range of typically medium size ponds and lakes. Area varied between 151m^2^ (N=11) for pools, 1113m^2^ (N=59) ponds, 6458m^2^ (N=27) for small lakes, 38687m^2^ (N=14) for medium size lakes, and 65169m^2^ (N=3) for large lakes. The area is within the boundaries of Pyrenees National Park, France and comprises a series of postglacial catchments. Catchment geology varied between the various valleys and it was dominated by two large geologic units: in the central area and at the extreme east, lake catchments lie on acidic bedrock (granite batholith) while in between, granitic batholiths were surrounded by metasedimentary and sedimentary materials such as slate, limestone and sandstone (Zaharescu, 2011).

**Fig. 1.**
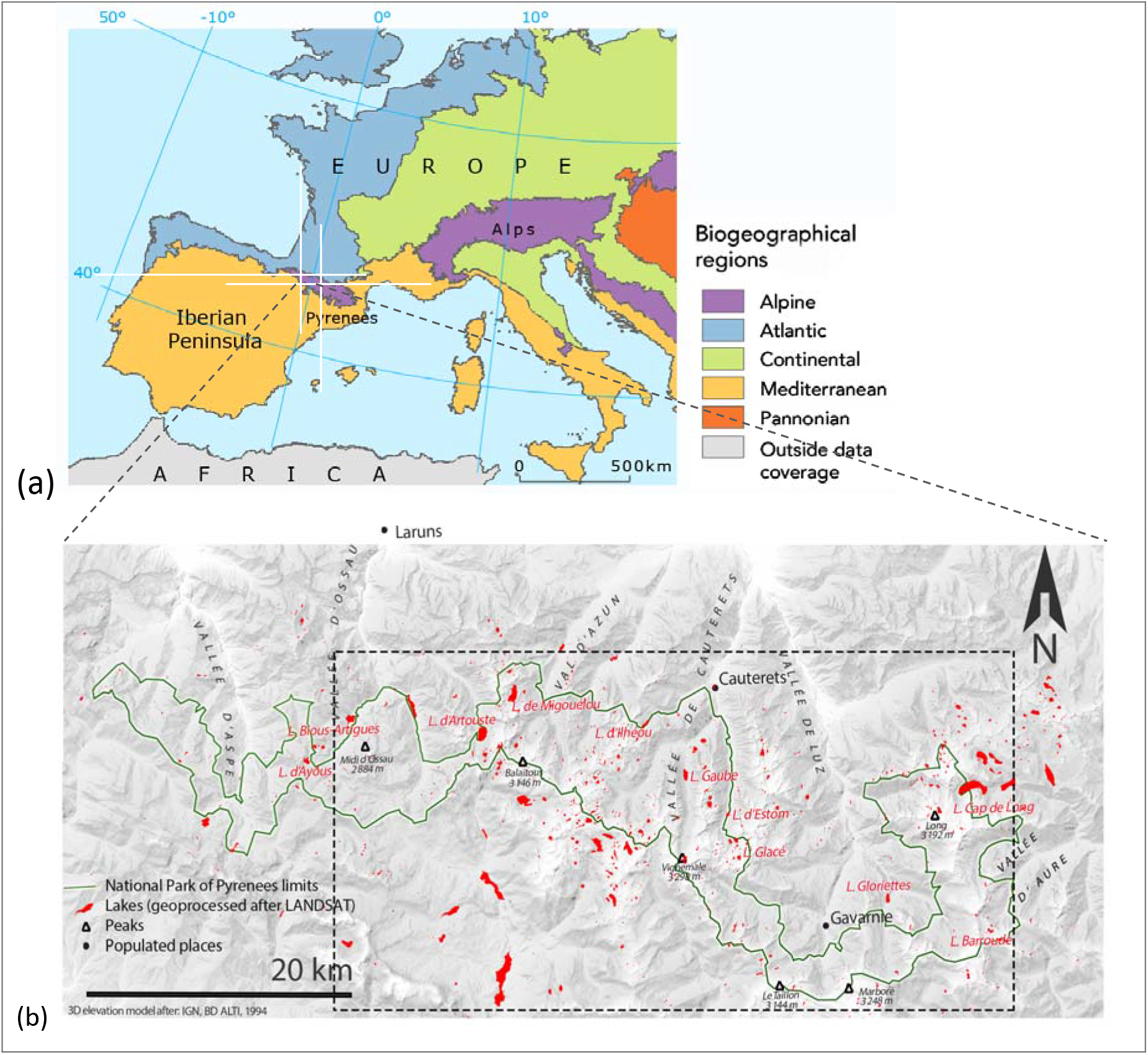
(a) Major biogeographical regions of Europe (after EEA, 2001). (b) High altitude lakes distribution in the Pyrenees National Park, France (green boundaries). Only lakes within park boundaries, which are enclosed in the dash line box were considered for this study.

Most of the study lakes are above the tree line (altitudes ranged from 1580-2501 m a.s.l.; mean=2214m a.s.l.), and they are largely undisturbed by human activity. Pastoralism, leisure fishing and trekking are among the very few activities allowed in the park. Only one sampled lake was transformed into reservoirs (Lake Ossoue), and it is being used as freshwater reserve. The great majority of study lakes/ponds are oligotrophic. Their proximal catchment area (roughly 10-20 m around the lake) has generally low vegetation coverage (<20%), but this varies according to topography and location. Loose rocks dominate on most of the lake shores, though they were more abundant on the steeply slopes of granitic catchments (Zaharescu, 2011).

The hydrological network, consisting of temporary and permanent lakes, ponds, pools and streams, is a natural legacy of the last glaciers retreat more than 5000 years ago. Water input in most lakes is by direct precipitation and permanent streams; glaciers and springs were present only in a few cases. Surface connectivity between lakes varied for the lakes investigated. Slope/bank snow coverage at the time of sampling was generally low, but had generally higher coverage at the head of catchments. Water pH was generally neutral (mean = 7.59) but varied between 5.2 and 8.8. Conductivity was also variable, ranging between 2 and 267 μS cm^-1^ (mean= 40μS cm^-1^).

### 2.2 Sampling strategy and data collection

An exhaustive assessment was conducted for each visited water-body (Fig. 1). It included littoral macroinvertebrates, water pH and conductivity, the presence of vertebrate predators, i.e. frogs and trout, ecotope properties of near catchment, riparian vegetation assemblage and geolocation. We use the term “ecotope” to denote the integrated physical elements of a landscape that underlie an ecosystem, and that exchanges matter and energy with the surrounding environment.

Macroinvertebrate sampling deliberately targeted the littoral zone. This area generally supports far larger and more diverse populations of benthic invertebrates than the deeper zone (Vadeboncoeur et al., 2011). The littoral is also likely to relate more directly to the nearby riparian and catchment factors. Semi-quantitative 3-minute kick-samples were collected in each lake using a standard pond net (Frost et al., 1971). Samples were collected at short distances while moving around the lake perimeter to cover different micro-habitats in proportion to their occurrence. Littoral substrata was highly variable and ranged from boulders to fine sands, vascular plants, mosses and algae. A composite sample (3-10 subsamples) was collected at each lake. Due to size and challenging topography almost half of the perimeter was sampled in each lake. All substratum types (rocks, cobbles, coarse and fine sand, epilithic moss, etc.) were sampled down to 60 cm water depth. Subsequently all samples were preserved in 96% alcohol for a comprehensive laboratory sorting and analysis. Benthic organisms were identified down to the lowest possible taxonomic level using Tachet et al. (2002) key, and counted under a stereomicroscope. The lowest taxonomic level identified (down to genus and species in some cases) of living and subfossil taxa will be regarded as morphotypes henceforth. For most statistical tests a family/subfamily level resolution was used (Supplementary List 2).

Additionally, water pH and conductivity were recorded at the surface and the bottom (± 5m off the shore) at each site with portable pH/conductivity probes. The water was collected with a standard bottom water sampler, following a clean protocol (Zaharescu et al., 2009). Presence of frogs (Rana temporaria) was visually inspected at each site. Trout presence data at each location was obtained from the stocking records maintained by the Pyrenees National Park. Furthermore, at each location a number of landscape factors were visually approximated according to dominant units. They were: nature of water input and output, tributary discharge, water-body size, % vegetation covering slopes and shore, slope, geology, presence of aquatic vegetation, shore development (fractal level), presence of snow deposits on the shore and in the catchment (%), catchment type and surface connectivity with other waterbodies (Zaharescu, 2011, Zaharescu et al., 2015a).

Riparian vegetation composition (presence/absence data) was recorded down to species level in the field at each site (for 50-100 % of lake perimeter), or on plants collected in a vasculum and identified off site, using multiple identification keys (Grey-Wilson and Blamey, 1979; Fitter et al., 1984; and García-Rollán, 1985). A detail description of the procedure is described in Zaharescu (2011) and Zaharescu et al., 2015b. Lakes geographical position was recorded with a portable GPS device and provided in Supplementary List 2.

### 2.3 Data analyses

Statistical data analyses included principal component analysis (PCA), fuzzy set ordination (FSO), multidimensional FSO (MFSO), cluster and indicator species analyses. For this, environmental factors were split into groups, i.e. geolocation, landscape/ecotope, predation, water general chemistry and riparian vegetation.

#### 2.3.1 Principal Component Analysis

First, the landscape variables were reduced to a limited number of meaningful composite factors (Principal Components) by using the PC regression scores from PCA, after maximizing their fit to variable groups (Varimax rotation). These composite factors were used as predictors of littoral zoobenthos in further fuzzy set analysis (Table 1). By default, the principal components of PCA with this rotation are uncorrelated.

**Table 1.**
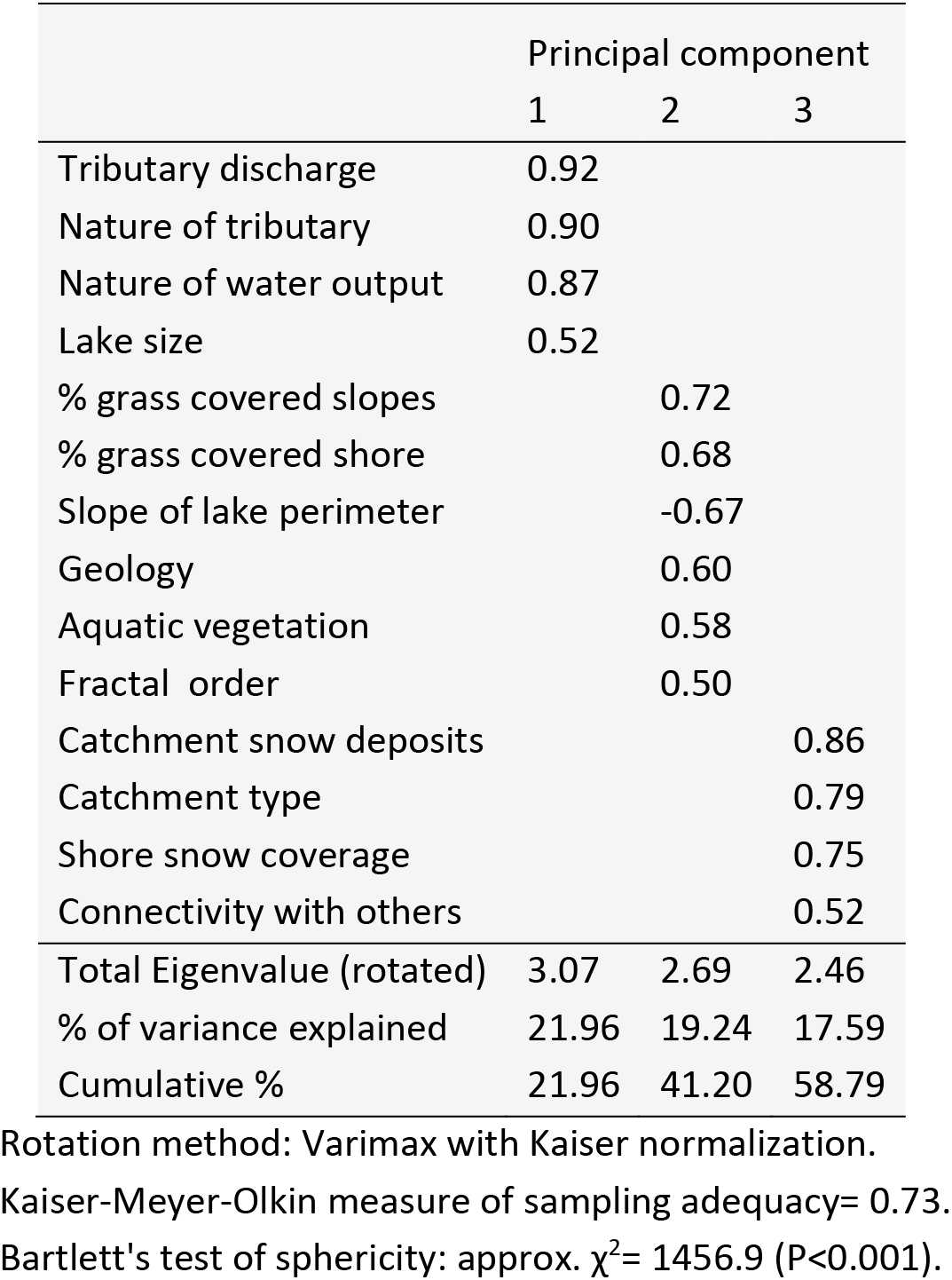
Association between catchment variables characterising the Pyrenees lakes, and PCA components. Only highest variable correlation with any of the components is displayed. This allowed interpret PC1 as hydrodynamics, PC2 as geo-morphology and PC3 as topography formation.

|  | Principal component |  |  |
| --- | --- | --- | --- |
|  | 1 | 2 | 3 |
| Tributary discharge | 0.92 |  |  |
| Nature of tributary | 0.90 |  |  |
| Nature of water output | 0.87 |  |  |
| Lake size | 0.52 |  |  |
| % grass covered slopes |  | 0.72 |  |
| % grass covered shore |  | 0.68 |  |
| Slope of lake perimeter |  | -0.67 |  |
| Geology |  | 0.60 |  |
| Aquatic vegetation |  | 0.58 |  |
| Fractal order |  | 0.50 |  |
| Catchment snow deposits |  |  | 0.86 |
| Catchment type |  |  | 0.79 |
| Shore snow coverage |  |  | 0.75 |
| Connectivity with others |  |  | 0.52 |
| Total Eigenvalue (rotated) | 3.07 | 2.69 | 2.46 |
| % of variance explained | 21.96 | 19.24 | 17.59 |
| Cumulative % | 21.96 | 41.20 | 58.79 |

Rotation method: Varimax with Kaiser normalization.
Kaiser-Meyer-Olkin measure of sampling adequacy= 0.73.
Bartlett's test of sphericity: approx. $\chi^2= 1456.9$ ( $P<0.001$ ).

#### 2.3.2 (Multidimensional) Fuzzy Set Ordination

To analyse the relationship between littoral zoobenthos composition (presence/absence data) and environmental gradients we used fuzzy set ordination (FSO) followed by stepwise multidimensional FSO (MFSO; Roberts, 2008). For these a distance (dissimilarity) matrix computed with Sørensen similarity index of invertebrate presence/absence data was first calculated. This gave a measure of similarity between sites based solely on biotic composition (Boyce, 2008). Additionally, two more variables assumed to describe zoobenthos community structure were used in a (M)FSO with vegetation presence/absence data matrix (Sørensen similarity index). They were taxon (family) richness and sequential diversity comparison index, which is a simplified method for estimating relative differences in biological diversity (SCI; Barbour et al., 1999), and allowed considering morphotypes in the analysis (Equation 1). *Run* describes the morphotype and *taxon*, the family

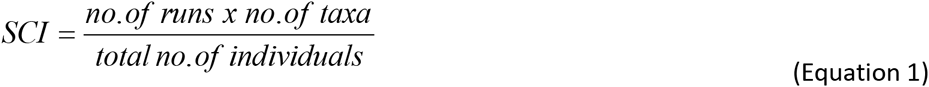

Fuzzy set ordination (FSO) concept (Roberts, 1986) is a generalised alternative to traditional ordination approaches-such as canonical correspondence analysis, in which cases are assigned gradual membership (fuzzy) values ranging from 0 to 1 (Roberts, 2008), instead of 0 or 1 (i.e. in-or-out of a given set) like in classical statistics. FSO is expected to perform better than other models on more complex data sets, and it is insensitive to noise in environmental factors and rare species (Roberts, 2009).

Variables were first screened in turn in FSO, and those with highest correlation with the zoobenthos distance matrix (at >95% efficiency) were retained for further MFSO. Technically, in MFSO, a FSO is performed on the variable that accounts for most of the variation first. Then, the residuals of the analysis are used with the next most important variable. The process is repeated until no more variables are left. Because only the fractions of variable membership that are uncorrelated are used by MFSO, each variable selected by the model is regarded as an independent process. This gives a high interpretability to the model (Roberts, 2008). Visually, the effect extent of each variable can be assessed by the increment in the correlation value attributable to that variable.

A total of 1000 random permutations were subsequently performed to test the significance of each variable in FSO/MFSO. Where the distance matrix was disconnected (sites/groups of sites with no shared species) or the dissimilarity was too high, a step-across function was applied to improve the MFSO. This finds the shortest paths to connect groups and removes rare observations/ groups of observations (Oksanen, 2008).

Because trout and frog variables were binary, and to achieve more accurate R^2^ in the model, these variables were standardized by Hellinger transformation (Legendre and Gallagher, 2001) before using them in FSO.

#### 2.3.3 Mantel test

To further assess the potential effect of riparian vegetation composition on major littoral invertebrate composition a Mantel test was performed on their distance matrixes. These matrixes were calculated with Baroni-Urbani & Buser similarity index. This index was preferred as it maximises the Pearson product-moment correlation coefficient between the two matrixes. A high significance of the correlation procedure was drawn after 9999 random permutations of Monte Carlo test. Mantel test was further used to test for the relationship between vegetation structure (computed using Sorensen similarity index) and zoobenthos family richness and morphotype diversity.

#### 2.3.4 Community analysis

Finally, the littoral zoobenthos data (family presence/absence) was analysed for co-occurring taxa and their ecotope preferences. This was achieved by clustering the sites on the basis of shared species, and applying indicator species analysis for each resulting cluster. First, a flexible linkage Pair-Group Method using the Arithmetic Averages (PGMA; method parameter = 0.85) cluster analysis was run on a distance matrix computed from Sørensen similarity matrix of families presence/absence data. Plotting cluster solutions in discriminating space (by discriminant analysis) helped evaluate the reliability of cluster solution. Secondly, indicator species analysis was run at the nodes of the major clusters to identify invertebrate families that represent the resulting lake groups.

FSO and LabDSV packages were used to compute FSO and MFSO (Roberts, 2007a; Roberts, 2007b); ADE4, CLUSTER and FPC packages for Mantel test, clustering (Thioulouse et al., 1997; Kaufman and Rousseeuw, 1990; Hennig, 2005), and LabDSV for indicator species analysis (Dufrene and Legendre, 1997), all for the R statistical language and environment (R Development Core Team, 2005).

## 3 Results and discussion

### 3.1 Littoral diversity, landscape structure and scale

#### 3.1.1 Large geographical gradients

Biome variability across large geographic areas generally follows large-scale gradients of climate and topography. Understanding how littoral ecosystem responds to these gradients is important in predicting effects from global environmental changes. Results of FSO/MFSO of family composition against altitude, latitude and longitude showed that individually, the three factors could reliably predict littoral taxa composition (Fig. 2). The relative contribution of these variables to MFSO and their cumulative value are illustrated in Fig. 3. Longitude exerted by far the largest independent contribution, while altitude and latitude appeared to incorporate a large covariant component with the former, as shown by their low significance, P, as independent factors.

**Fig. 2.**
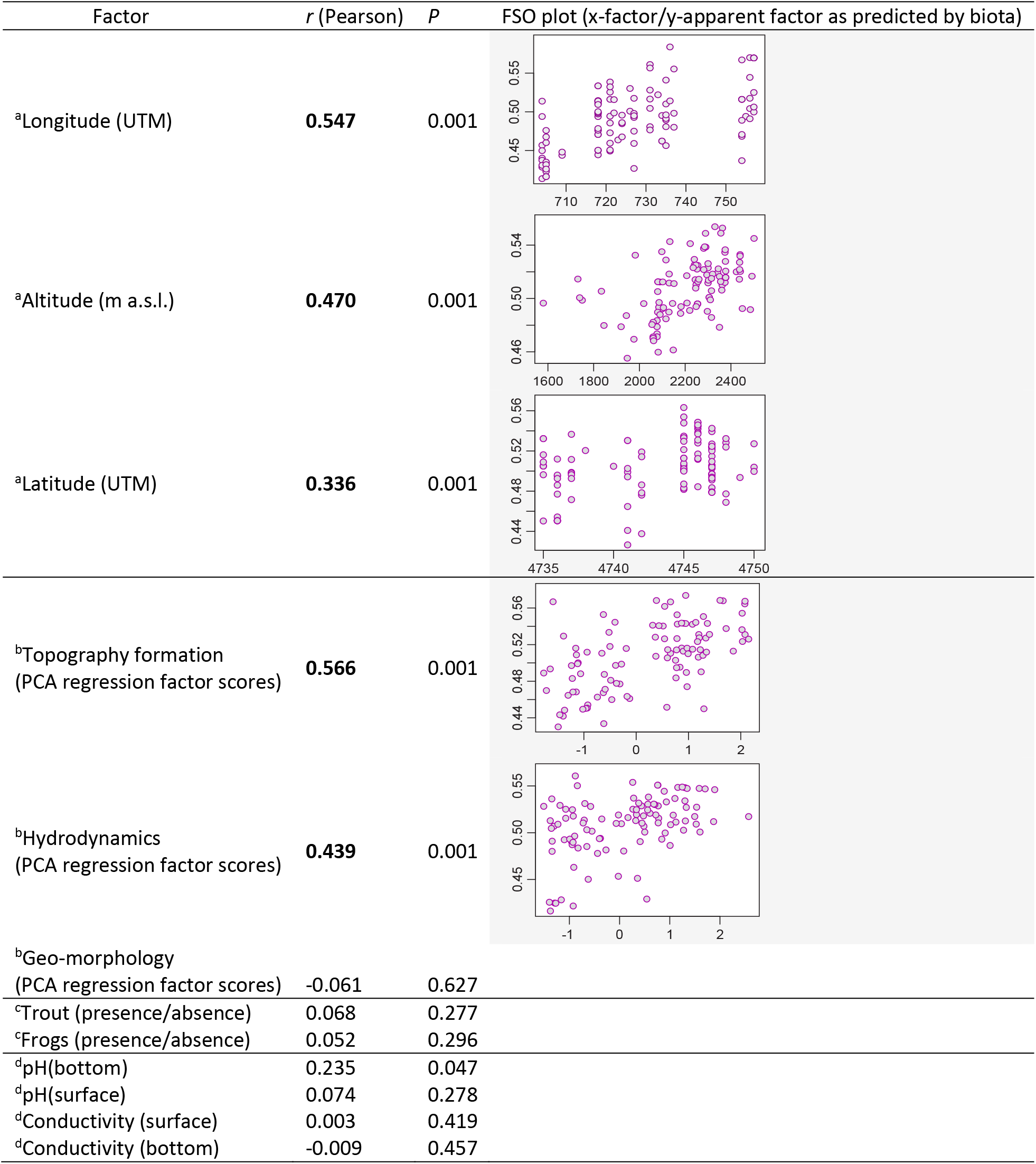
One-dimensional fuzzy set ordination (FSO), showing the response of littoral invertebrate family structure to environmental variables in the Pyrenean lakes. Indices represent: (a) geolocation, (b) composite catchment (Table 1), (c) predation and (d) water physico-chemistry. Correlations are listed in descending order. Variables with highest influence in the model (correlations >0.3, in bold), also shown in plots, were retained for multidimensional FSO. *P* represents the probability. Predation variables were Hellinger transformed (Legendre & Gallagher, 2001) previously to being used as predictors in the analysis.

**Fig. 3.**
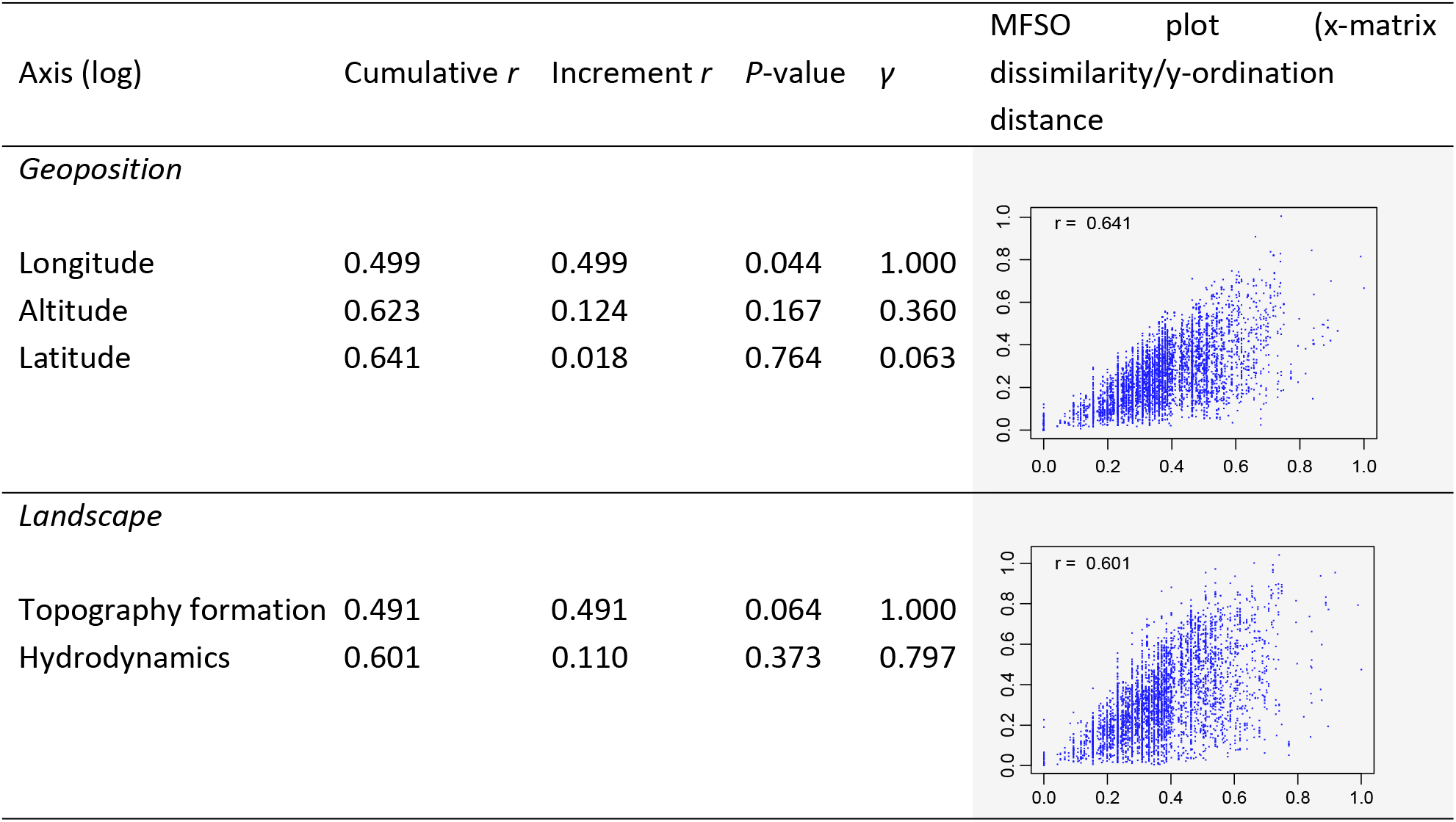
Multidimensional response of major littoral invertebrate composition to geoposition and composite catchment factors in multidimensional FSO (MFSO) with step-across improvement. Variables are added to the model as log transformed, in the order of their decreasing fuzzy correlation (Pearson) with biota dissimilarity matrix. Permutation number = 1000. *γ* (gamma) represents a vector of the fraction of variance of a factor that is independent of all previous factors. Due to the high-dimensional variability of the dissimilarity matrix, the correlation probability for the one-dimensional solution sometimes has low significance, but it is still valid.

Compositional/functional changes in zoobenthos across large horizontal and vertical gradients have been reported before, and whole biome models have been used to evaluate changes in taxon distribution likely to occur with a changing climate (Colwell et al., 2008, IPCC, 2014). At an estimated 60km longitudinal span, the study area is relatively narrow. Nevertheless, latitude dominance in the model appears to be given by the area’ strategic position at the confluence of major biogeographic regions Atlantic and Mediterranean (Fig. 1), which experienced sharp ecosystem changes. This highlights the capacity of high altitude littoral ecosystem to reflect transitional gradients between major biogeographical regions over relatively narrow spaces, and this can potentially experience major climate change consequences.

#### 3.1.2 Catchment structure

Principal component analysis (PCA) revealed three composite factors (Table 1). These factors were interpreted as: PC1, hydrodynamics (summarising input size, input and output nature, and lake size); PC2, geo-morphology (i.e. % vegetated shores and slopes, shore slope, geology, aquatic vegetation and shore development); and PC3, topography formation (catchment type, % shore and catchment snow coverage, connectivity with other lakes). They are exhaustively reported in Zaharescu et al. (2015a). The response of littoral invertebrates to these catchment factors is illustrated in Figures 2 (FSO) and 3 (MFSO). Both, univariate and multivariate solutions of the model show that topography was the most important predictor of littoral biota composition at a high degree of confidence (p<0.06), followed by hydrodynamics (Fig. 2 and 3). Topography exerts its influence mainly through its structural variables: catchment type, shore and catchment snow coverage and connectivity with other lakes. These variables would sustain habitats at larger scale (e.g lake’s proximal catchment), and allow connectivity among populations of benthic communities, which need adequate habitats in both, aquatic and riparian areas for survival. For instance, lakes/ponds at the head of glacial valleys, with snow presence most of the year, would harbour functional taxa with adaptation for cryal environment, very low nutrient input, and short reproductive time. On the other hand, valley floor lakes would harbour organisms with longer emergence periods, requiring additional nutrient and material inputs from the catchment, and allowing more diverse periphyton communities that serve as food and microhabitats. This ecosystem would also likely be more vulnerable to larger periods of snow presence.

While hydrodynamics was significant in FSO (Fig. 2), its relatively small influence in MFSO is due high co-variabile effect with topography (Fig. 3). The secondary effect of lake hydrodynamics suggests contributions from water source and lake area. For instance, large stream-fed lakes generally maintain a continuous surface flow throughout the summer, would also maintain a generally low temperature and a heterogeneous structure of littoral habitats. Conversely, in relatively smaller waterbodies, dominated by catchment runoff and/or snow melt (therefore not sourced by continuous streams), the extent of littoral surface can vary seasonally and warm faster. These different ecotopes will allow the persistence of functional groups adapted to the distinct lake environments, and they will vary in concert with topography formation. This is supported by the results of studies conducted in other altitude environments, which found clear differences in biotic assemblages in spring-fed streams under different flow regimes (Danehy and Bilby, 2009).

#### 3.1.3 Effect of riparian vegetation

Many of the benthic invertebrates, particularly insects, also present terrestrial phases. The relationship between littoral and riparian ecosystems may therefore go beyond their simple proximity or nutrient provision. (M)FSO model found a significant effect of plant species composition on the invertebrate diversity and family richness (cumulative r=0.48, p<0.05; Fig. 4). Relatively low but significant relationship was also found between the compositions of vegetation and benthic invertebrates (Mantel test, Monte Carlo r= 0.16, p<0.01), which means commonly associated invertebrate groups are supported by commonly associated plant species. Although spatial covariability of flora and fauna along environmental gradients is not excluded, this relationship is meaningful in the sense that in the restricting alpine environment plant consortiums could provide niche separation for various competing invertebrates, including the terrestrial phases of most aquatic insects. This could include supplying nutrient for functional feeding groups, casing materials, microhabitats during short summer periods, and protection against excessive solar radiation (Gregory et al., 1991; Dudgeon, 2009). Other studies have highlighted the importance of riparian plant coverage to macroinvertebrate communities along streams, especially in strong transitional gradients such as grassland-forest (Stone et al., 2005), but also the vegetation type (Cummins et al., 1989; Angradi et al., 2001). Our findings support the idea that sparsely vegetated altitude catchments provide important functional links between riparian vegetation composition and the diversity, richness and functional composition of benthic invertebrates.

**Fig. 4.**
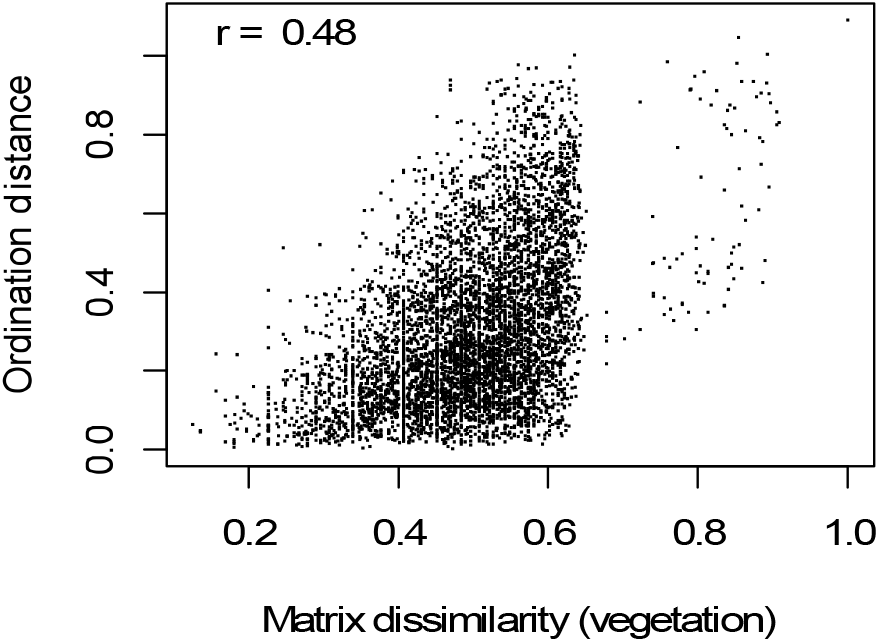
Relationship between riparian vegetation structure and littoral invertebrate morphotype diversity and family richness in a bidimensional FSO. A step-across function improved the ordination. Number of permutations = 1000.

#### 3.1.4 Vertebrate predation and major water chemistry

Littoral productivity is vital for supporting higher trophic levels in lakes (Vadeboncoeur et al., 2011), and the presence of predators such as fish or amphibians, particularly in alpine lakes can result in a top-down driven ecosystem (Eriksson et al., 1980). Results of the relationship between the presence of fish and amphibians, and invertebrate groups surprisingly showed no effect (Fig. 2). This is evidence that the broad composition of littoral fauna is highly resilient to vertebrate predation. It is possible that predators were size selective, affecting the abundance of easily accessible groups, such as chironomids (Orthocladiinae and Chironominae) and planktonic crustaceans (Kernan et al., 2009; Syväranta and Jones, 2009; Schilling et al., 2009). Or, a coarse littoral substrata and protection mechanisms insects use in alpine lakes could be good defence mechanisms against vertebrate predation. Niche segregation between aquatic and terrestrial environments could have also played a role. It is known that alpine lake frogs would largely prey on the more abundant terrestrial (adult) insects (Vieites et al., 1997), which helps them maximise nutrient sequestration during the short period these lakes are unfrozen. Carlisle and Hawkins (1998) who observed that physical habitat might be more important than predation in structuring benthic communities in trout-stocked mountain lakes further support our results. Clearly, these interesting results merit further evaluation.

Water pH and conductivity, measures of acidity and total ionic/nutrient content – important lake parameters, could not explain diversity variation in major zoobenthic groups (Fig. 2). They are both major indicators of bedrock geology/weathering and lake metabolism, and can change significantly during thaw periods in mountain lakes, influencing biotic composition (Olofsson et al., 1995). The low relationship observed for either surface or lake bottom, suggests that their natural/seasonal variability in each lake may be strong enough to offset a direct response from biotic communities at a broader scales.

### 3.2 Major littoral communities

Harsh environment and low nutrient of headwater habitats present unique challenges to littoral biota. Strong biogeographical fragmentation is therefore expected, which could result in communities that are strongly dependent to local habitats. Flexible hierarchical clustering and indicator taxa analyses identified three large lake groups hosting characteristic biota (Fig. 5 and Table 2).

**Fig. 5.**
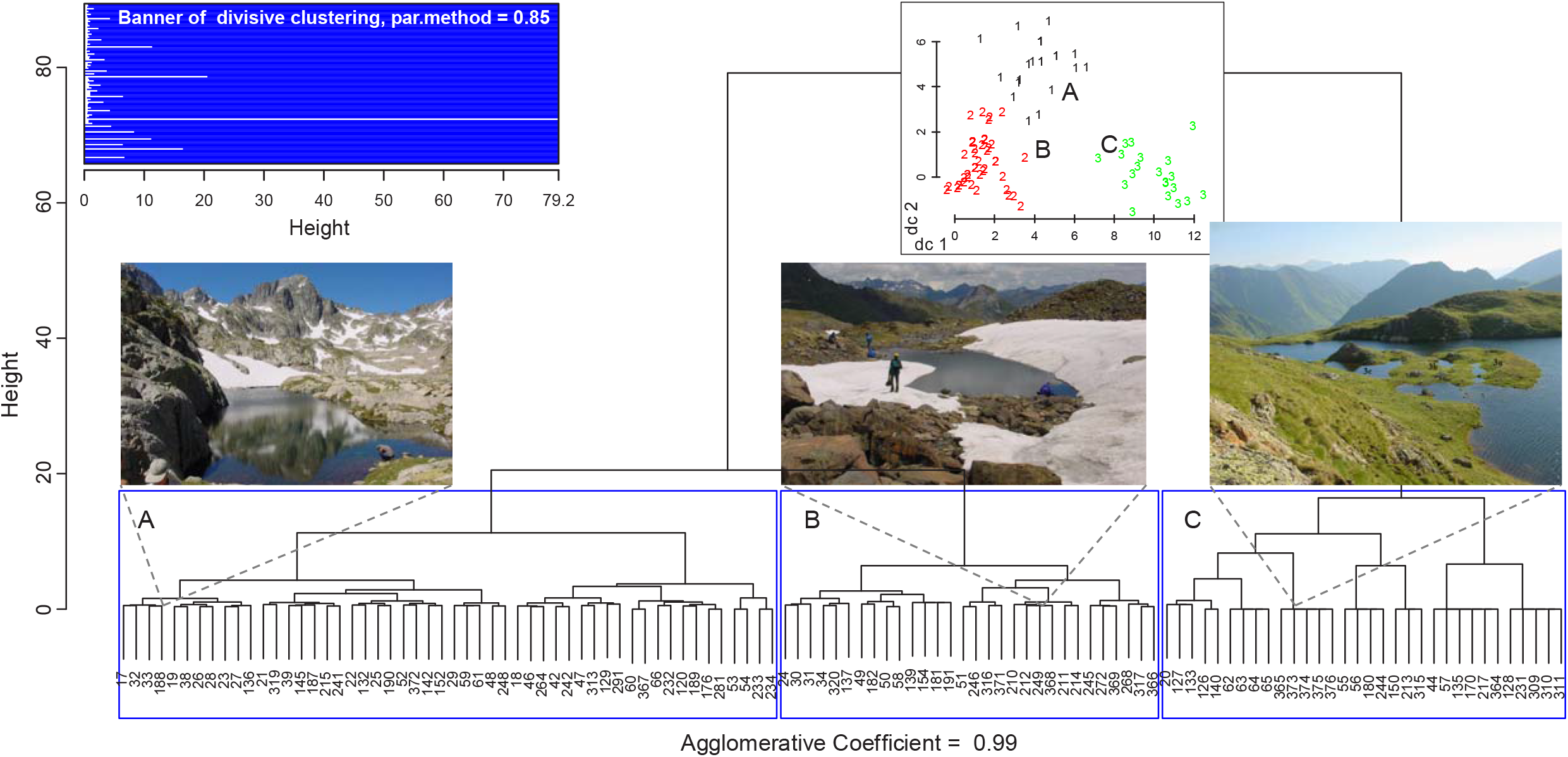
Major lake/ecosystem groups (A, B and C) as identified by hierarchical cluster analysis (flexible linkage, parameter = 0.85) based on shared littoral invertebrate families. A plot of cluster solutions in discriminating space (inset) demonstrate an effective clustering. Illustrated are: (A) Cambales Valley lake, (B) Mares de Montferrat, Ossoue Valley and (C), Barroude Petit, Aure Valley. The results are from an analysis of 114 lakes and 46 major invertebrate groups.

**Table 2.**
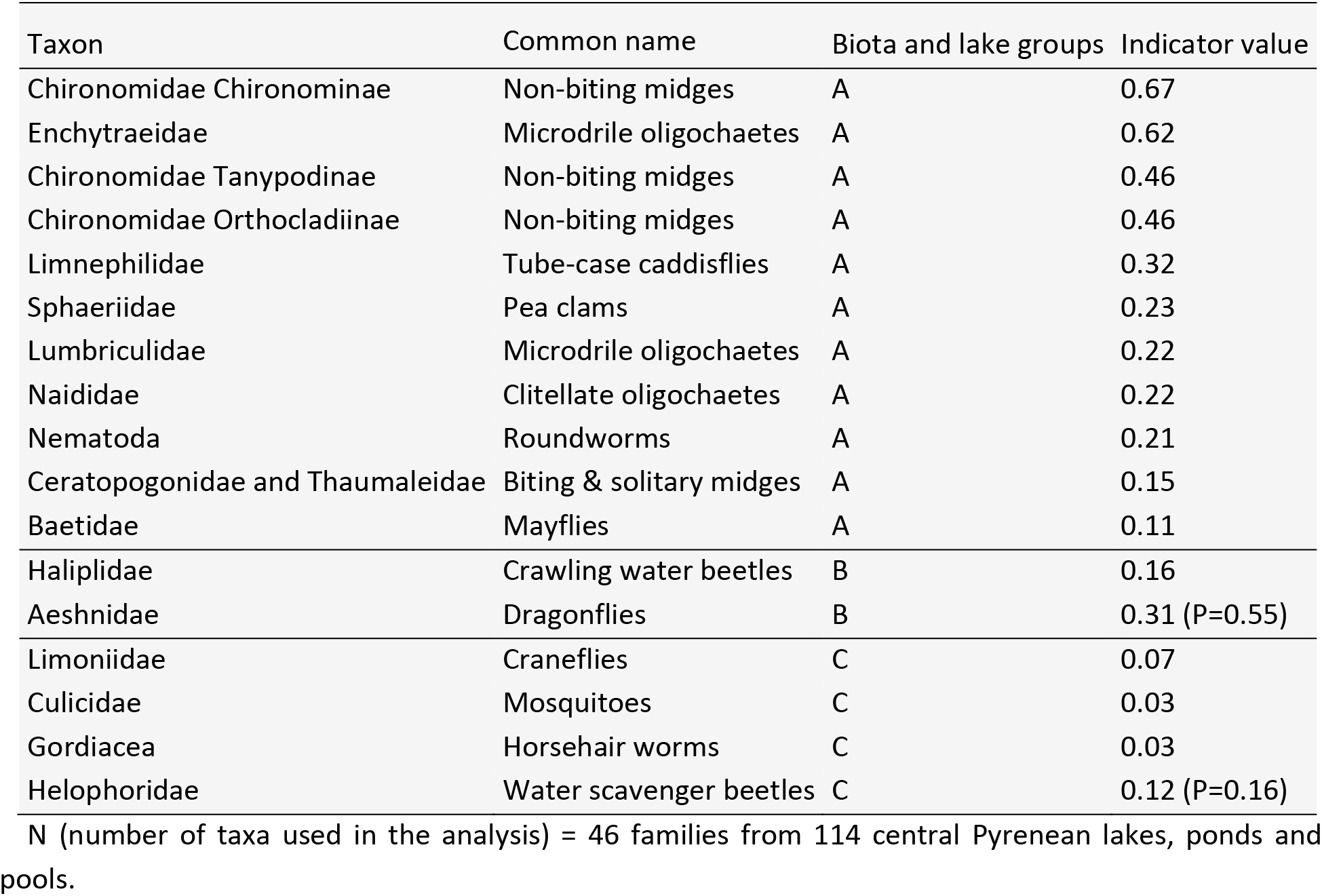
Zoobenthic communities with significant association to lake groups (from prior cluster analysis), as given by indicator taxa analysis. A subject was classified into a group for which the indicator value was higher and significant (i.e. strong preference). Significance level is <0.05, unless stated.

The first lake community (type A; Table 2) was the largest, and comprised a significant number of spring-dwellers, tolerant to wide ranges in temperature, altitude, flow regime, pH and micro-habitats (e.g. epi and endobenthic, rock surfaces and epiphytic). They were mostly of gill and tegumentary respiration, and a feeding strategy largely detritivorous and microphytes, but a small proportion were predatory (e.g. Tanypodinae larvae) and parasitic (nematodes). Their dispersion mode was mostly passive aquatic and aerial, which facilitates habitat connectivity (Tachet et al., 2002). The relatively wide ecological breadth of this group (eurytopic) means they can colonise a variety of headwaters. Association of Sphaeridae bivalves, Oligochaeta and Lumbriculidae worms with various members in this community has also been reported in headwaters of other regions, including the Oregon Coast Range and the Himalayas (Danehy and Bilby, 2009; Manca et al., 1998). The lakes group with this community are in Fig. 5.

Omnivorous beetles and predatory dragonflies represented the second community (type B). Both are strong flyers as adults, capable of active colonisation and maintaining connected populations not always at easy reach (Table 2). They have long life cycles (>1year) and tolerate a wide range of temperatures. They have affinity to low water flow regime and heterogeneous microhabitats (Tachet et al., 2002). Figure 5 displays lakes cluster sharing this littoral group.

The third littoral community (type C), had a low indicator value (Table 2). It was formed by craneflies, mosquitoes, water scavenger beetles and their parasitic worms. They share an aerial respiration (except gordiacea which are endoparasites in their larval stage) and a passive-to-active aerial dispersion mode in their adult stage. They tolerate a wide range of temperature and epibenthic microhabitats, with easy access to water surface where they breathe. Their feeding strategy was also diverse: shredders (Limoniidae), microphytes (Helophoridae) and, microinvertebrates and fine suspended matter (Culicidae) (Tachet et al., 2002). Females of most mosquitoes are ectoparasites.

Further boxplot comparisons revealed that these communities did not display distinct preferences along the assessed catchment-scale variables (not shown). This, together with the wide ecological tolerance revealed by their taxon composition suggests ubiquitous distributions, which may have resulted from natural evolution of lake ecosystems, or they were determined by lake/terrestrial factors beyond those analysed herein.

## 4 Conclusions

The findings simplify the complexity and highlight the level of connectivity between the littoral ecosystem and catchment heterogeneity (physical and ecological) in high altitude environment, at a wide range of scales. Littoral invertebrates responded significantly to large-scale horizontal and vertical gradients, dominated by longitude. The short E-W span ecosystem changes in the study area is consistent with a sharp transition between two major European climates: Mediterranean and Atlantic, which dominated the typical Holocenic altitudinal colonisation. The central Pyrenees are therefore a heterogeneous ecoregion in terms of their littoral ecosystemdistribution.

At catchment-scale, topography-related variables were the strongest predictors of littoral community composition, followed by hydro-dynamics. Topography controls habitats at relatively large scale, through traits such as lake basin morphology and riparian colonization. Hydro-dynamics had a secondary effect, suggesting that larger and smaller lakes host distinct littoral macroinvertebrate assemblages, likely associated to differences in water balance, biogeochemical nutrient fluxes in the catchment, and the general lake metabolism.

Locally, riparian vegetation composition affected littoral invertebrate community structure. Although generally poorly developed, different plant assemblages could provide distinct microhabitats for the terrestrial phases of aquatic insects, shelter against harsh conditions of solar radiation and wind, and weathered nutrients and casing materials for their aquatic phases. Aquatic vertebrate predation from trout and frogs, water pH and conductivity had no clear effect on the littoral zoobenthos composition.

Community analysis revealed three, relatively simple, functional associations of wide ecological tolerances, characterizing distinct ecosystems. They were of wide distribution across the studied region, a possible consequence of natural lake ecosystem evolution.

In the harsh climate and low nutrient alpine catchments, riparian and littoral areas are the most productive. The results demonstrate that the littoral ecosystem is connected in direct and indirect ways to a variety of physical, hydrological and ecological attributes from the terrestrial environment at scales extending from lake proximity, to its catchment and beyond. This is important as it helps appreciative the extent of terrestrial-aquatic interactions at high altitudes, and highlights their potential vulnerability to external stress from environmental changes. As climate-change continues to drive ecosystem shifts in the mountain biome, our results can serve as benchmark of relatively pristine systems. Our work also suggests that using the regional scale to characterize ecosystem processes in high altitude lakes is sufficiently robust, and can be confidently used in other studies on high altitude environments.

## Acknowledgements

This work was financially supported by Pyrenees National Park, France and Animal Anatomy Laboratory at Vigo University, Spain. We gratefully acknowledge field and data-support by: Andreea Vasiloiu, Catalin Tanase, Javier Fernandez-Fañanas, Nicolas Palanca-Castán, Jesús Giraldez-Moreira, Bruce Dudley and Cristina Castan-Lanaspa. We further thank Dave Roberts (Montana State University, USA) and Lasse Ruokolainen (Helsinki University, Finland) for the constructive conversation behind the statistical analyses.

## Authors contribution

Research and sampling campaign designs, A Palanca-Soler; data collection, DG Zaharescu, A Palanca-Soler and RN Lester; study design, sample and data analyses, DG Zaharescu and CI Burghelea; manuscript preparation, DG Zaharescu, PS Hooda and CI Burghelea.

## Supplementary Information

### Small lakes in big landscape: External drivers of littoral ecosystem in high elevation lakes

D.G. Zaharescu et al.

**Supplementary List 1.**
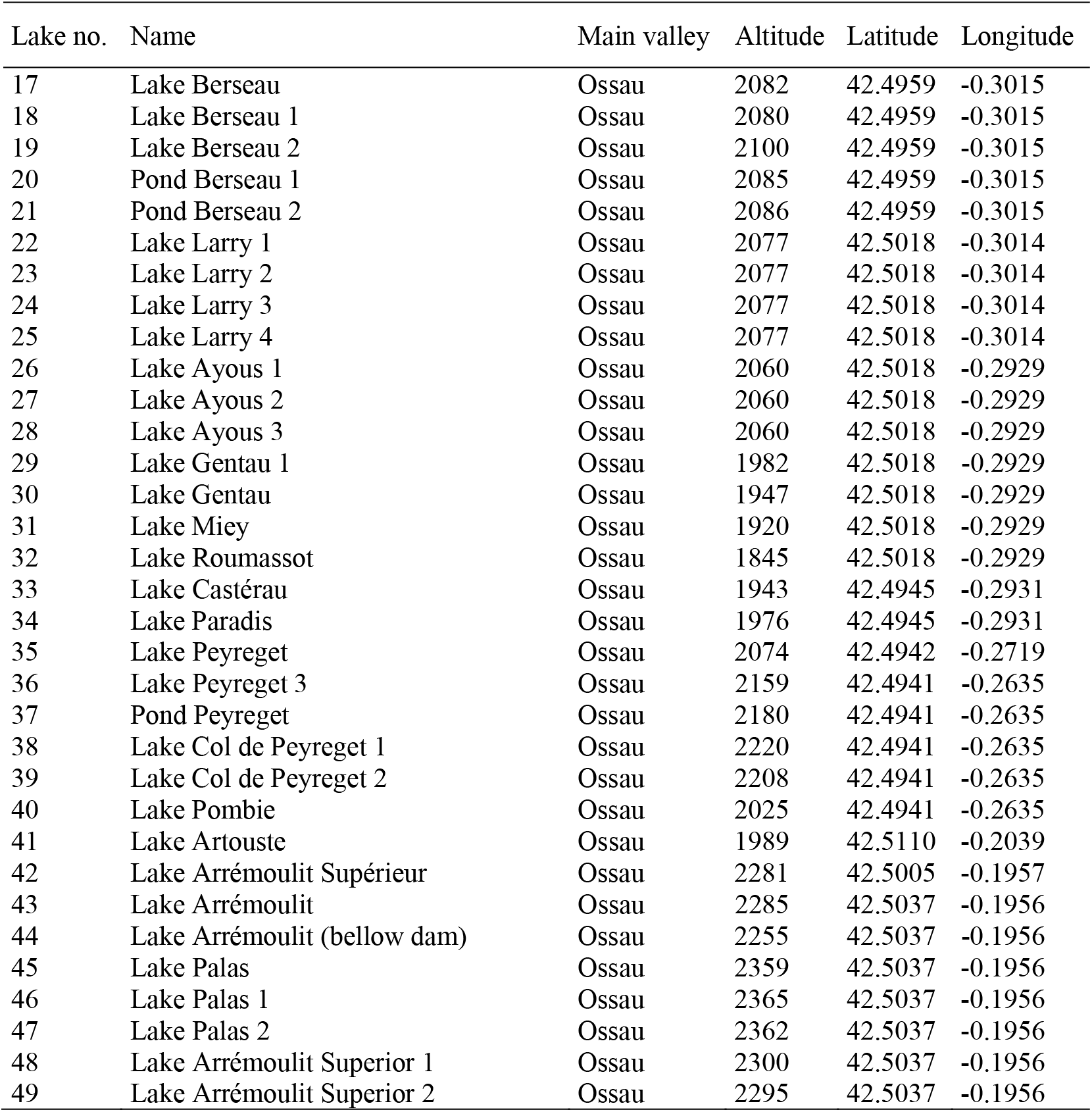
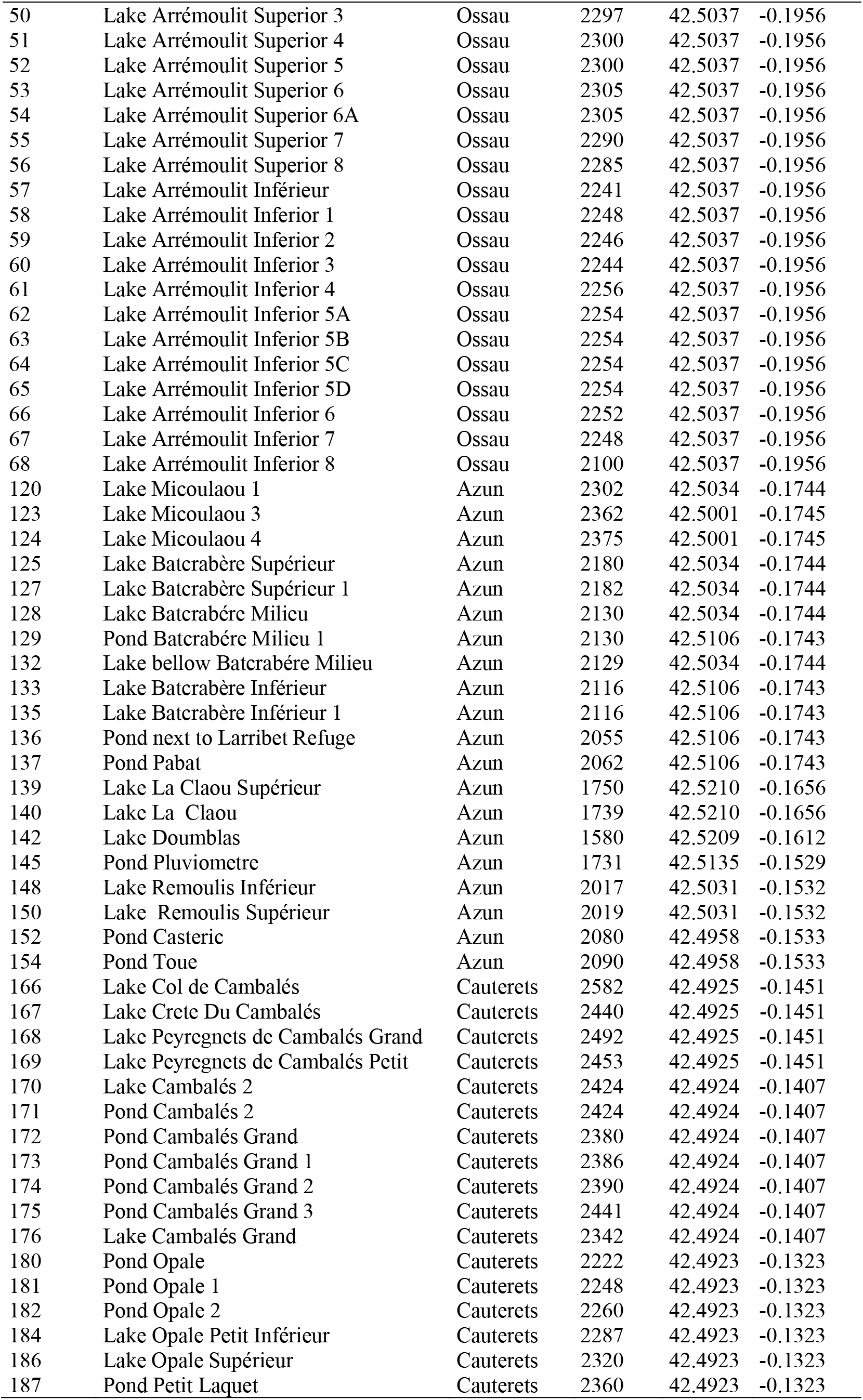
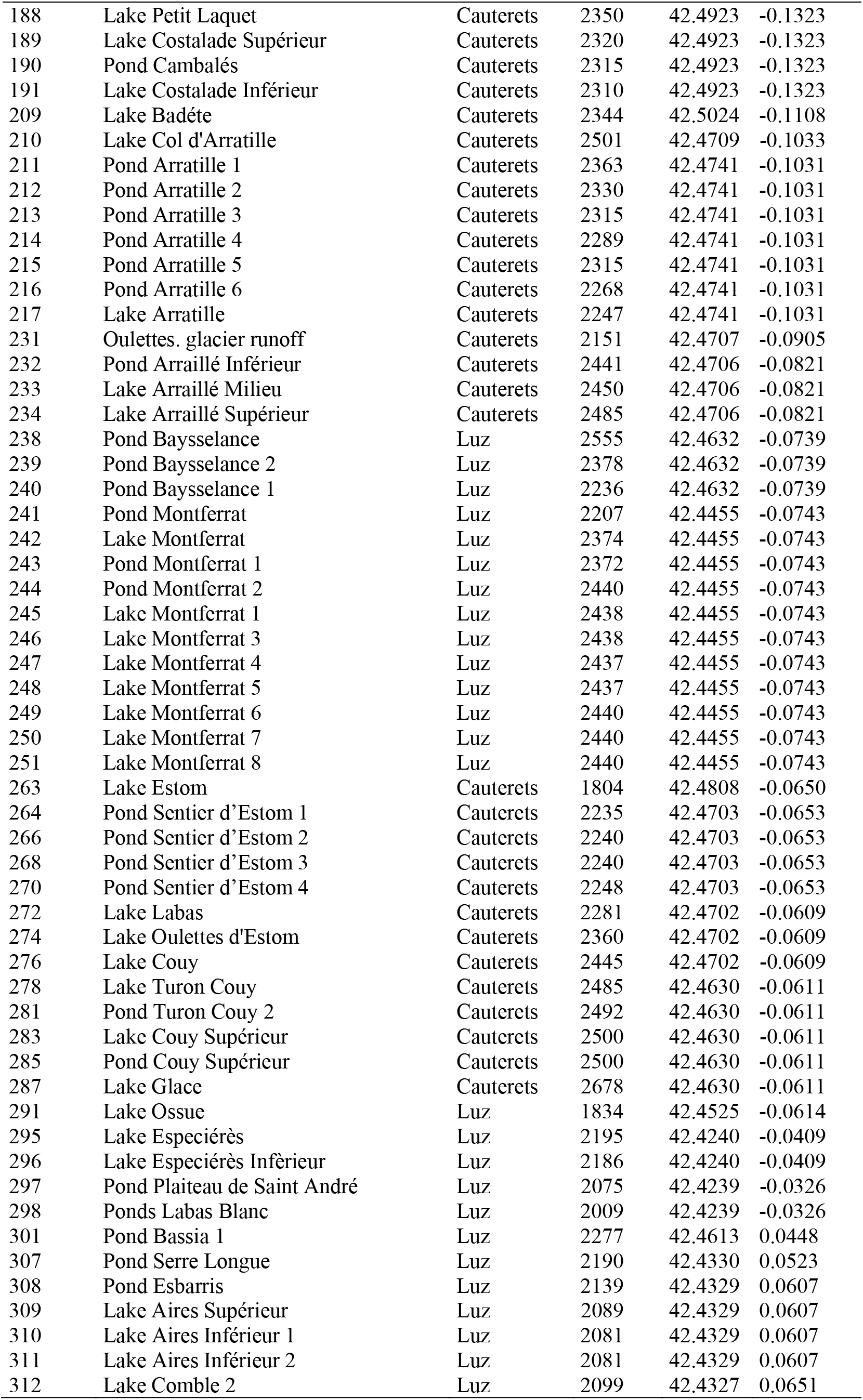
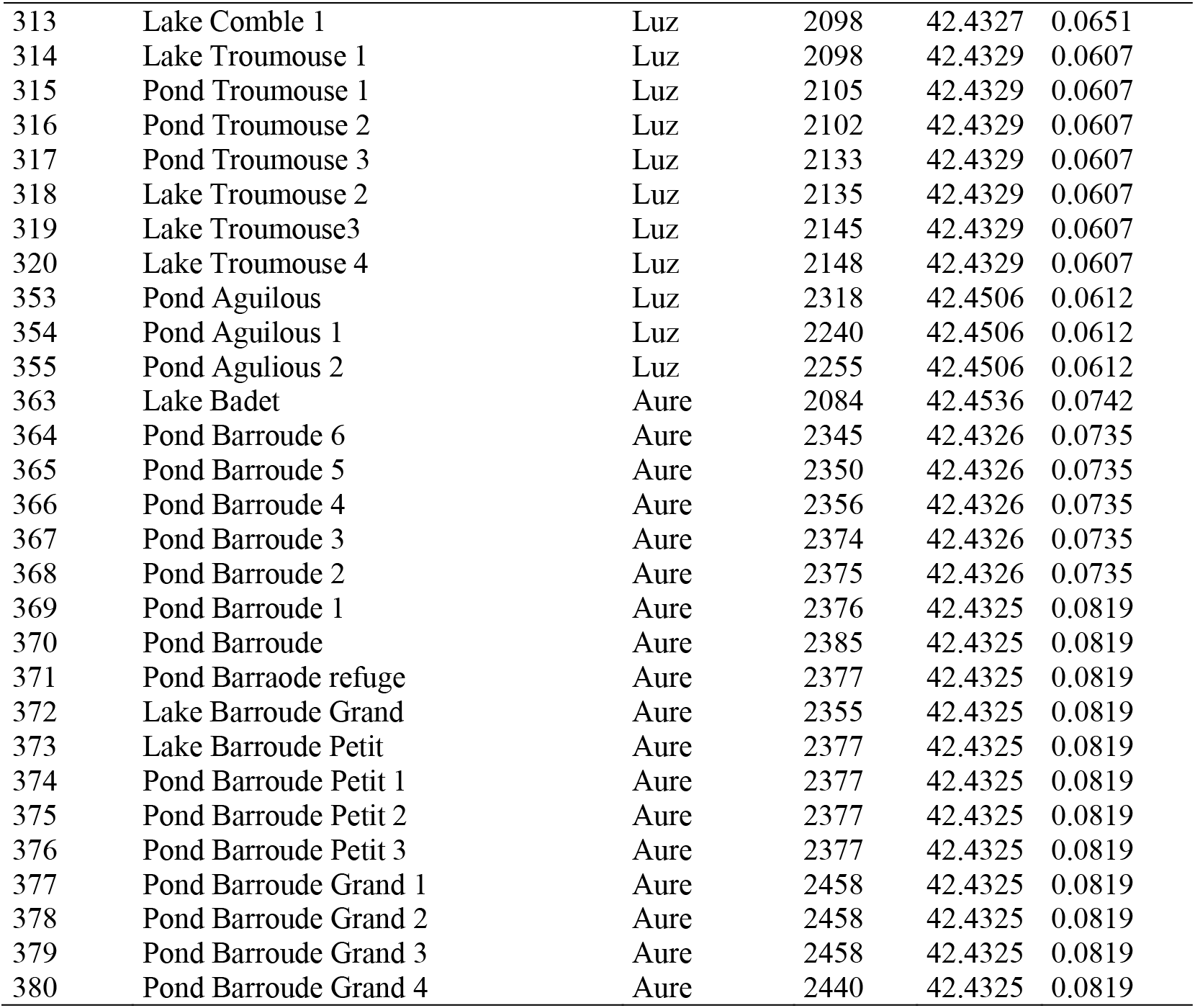
Lakes and ponds in the central Pyrenees National Park surveyed for this study, together with their main hydrographical network, altitude and geolocation (decimal degrees).

**Supplementary List 2.**
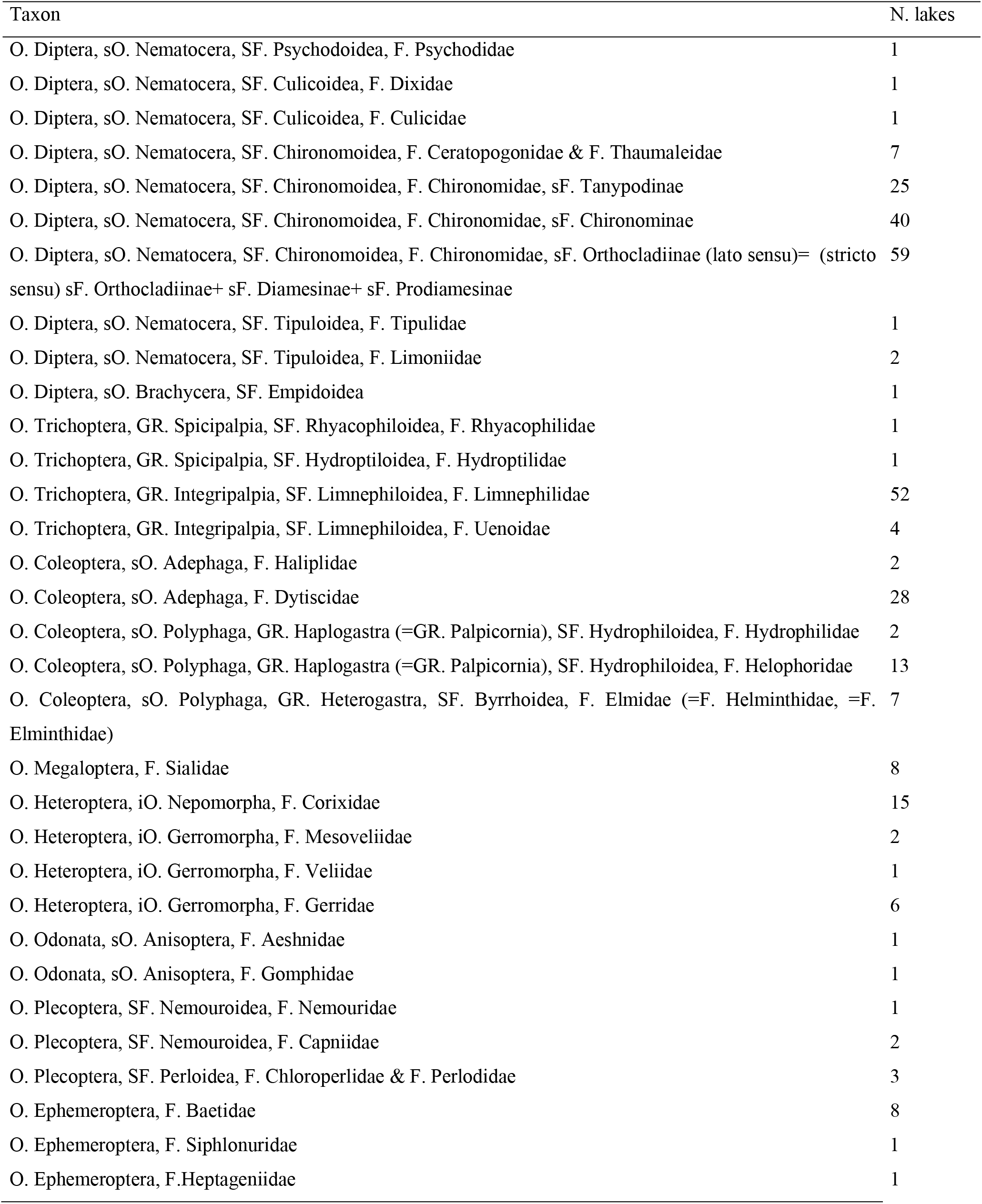
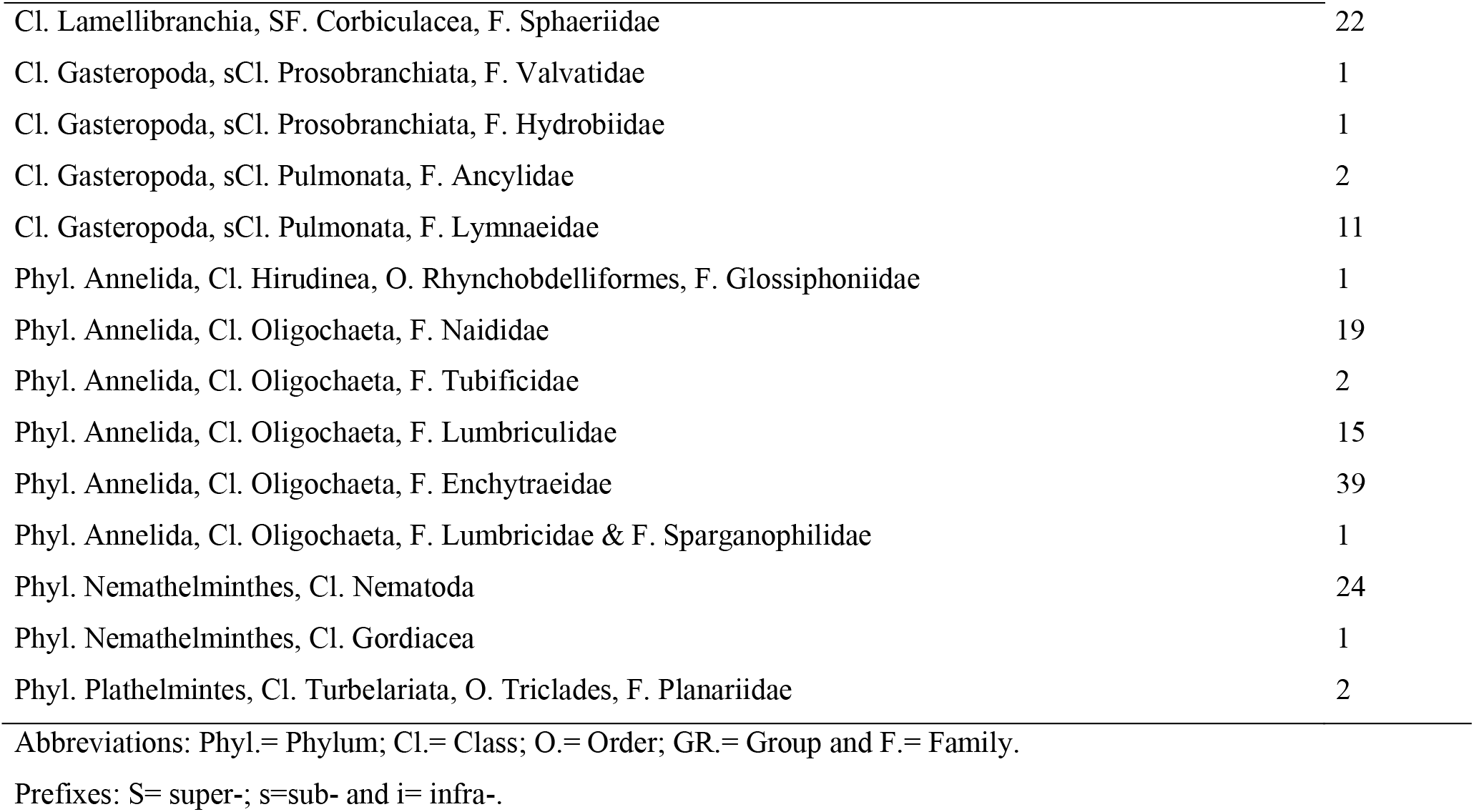
Major zoobenthos taxa and their incidence in the 114 lakes, ponds and pools of this study.

